# Spatial and seasonal dynamics of cholera (*Vibrio cholerae*) in an estuary in southern coastal Ecuador

**DOI:** 10.1101/110189

**Authors:** Sadie J. Ryan, Anna M. Stewart Ibarra, Eunice Ordóñez, Winnie Chu, Julia L. Finkelstein, Christine A. King, Luis E. Escobar, Christina Lupone, Froilan Heras, Erica Tauzer, Egan Waggoner, Tyler G. James, Washington B. Cárdenas, Mark Polhemus

## Abstract

Cholera emergence is strongly linked to local environmental and ecological context. The 1991-2004 pandemic emerged in Perú and spread north into Ecuador’s El Oro province, making this a key site for potential re-emergence. Machala, El Oro, is a port city of 250,000, near the Peruvian border. Many livelihoods depend on the estuarine system, from fishing for subsistence and trade, to domestic water use. In 2014, we conducted biweekly sampling for 10 months in five estuarine locations, across a gradient of human use, and ranging from inland to ocean. We measured water characteristics implicated in *V. cholerae* growth and persistence: pH, temperature, salinity, and algal concentration, and evaluated samples in five months for pathogenic and non-pathogenic *Vibrio cholerae*, by polymerase chain reaction (PCR). We found environmental persistence of strains O1 and O139, but no evidence for toxigene presence. *V. cholerae* presence was coupled to algal and salinity concentration, and sites exhibited considerable seasonal and spatial heterogeneity. This study indicates that environmental conditions in Machala are optimal for human cholera re-emergence, with risk peaking during September, and higher risk near urban periphery low-income communities. This highlights a need for surveillance of this coupled cholera– estuarine system to anticipate potential future outbreaks.

## Introduction

Cholera, a disease caused by the gram-negative bacteria *Vibrio cholerae*, remains a severe global threat to public health and development efforts (WHO 2013). A 2015 study estimated the global burden at 2.86 million cases, with 95,000 deaths, and 1.3 billion people at risk of cholera infection (Ali et al. 2015). An analysis of global cholera pandemics indicated that cholera outbreaks originate in coastal regions, often during flooding events, before spreading inland (Jutla et al. 2010). Studies suggest outbreaks of *Vibrio cholerae* can be explained by oceanographic variables (e.g., sea surface temperature, pH, salinity) and phytoplankton blooms, indicating the potential to predict disease outbreaks (Jutla et al. 2010; Jutla et al. 2013). Blue-green algae (BGA) has been described as a reservoir for environmental cholera (Islam et al. 2015), in part because its growth raises pH, encouraging *Vibrio* growth, but also as the major food source for zooplankton. The association between *V. cholerae* and zooplankton, wherein *Vibrio* are highly concentrated on their carapaces and internally, makes untreated surface water consumption in coupled zooplankton-phytoplankton hotspots a major potential point of infection (Lipp et al. 2002). Our previous work suggests that both predicted current and future coastal hotspots of cholera transmission are far larger than current epidemiological surveillance efforts can capture (Escobar et al. 2015). Indeed, the links between climate change and cholera brings the role of environmental *V. cholerae* persistence into sharp relief (Chowdhury et al. 2017), highlighting the importance of local surveillance efforts in vulnerable areas.

Estuarine systems are a natural intersection of coastal oceanographic conditions and human use; as productive systems for fisheries, port locations for transport, and rich riparian soils, they are a highly exposed interface. Coastal estuarine systems often represent subsistence-level dependence on the interface, in terms of artisanal fisheries, a higher likelihood of direct water use, and simply greater physical exposure by proximity. This renders them the most vulnerable of populations, most likely to be exposed to pathogens, and in the most flooding prone areas in the world (Nicholls 1995; Dixon et al. 2006; De Sherbinin et al. 2007; Hanson et al. 2011; Hallegatte et al. 2013; Cai et al. 2014).

The causative agent of cholera, *V. cholerae*, originates from estuarine waters. Evidence for this includes phylogenetic information and the physiological requirements for growth and persistence (Colwell and Huq 1994; Colwell 2004). *V. cholerae* is environmentally persistent in the Bay of Bengal (Bangladesh and India), along coastal areas in Latin America (Lipp et al. 2002; Mutreja et al. 2011), and in riverine, estuarine, and coastal waters around the world (Lipp et al. 2002). Cholera epidemics have historically followed coastlines (Colwell 1996), and *V. cholerae* can be transmitted to humans via a wide range of marine organisms, including zooplankton, aquatic plants, shellfish, and fish (Vezzulli et al. 2010).

Ecuador is a critical location to understand cholera and other climate and water-sensitive diseases due to its: (1) high potential for cholera outbreaks, and the high incidence of other climate-sensitive infectious diseases (e.g., leptospirosis, dengue), and (2) the strong influence of oceanographic conditions on local climate and flooding during El Niño events (Rossel et al. 1996; Rossel and Cadier 2009; Hanson et al. 2011; Hallegatte et al. 2013; Cai et al. 2014). In January 1991, cholera re-emerged in Latin America after more than a century without cases (Lacey 1995). In Ecuador, the 1991 cholera epidemic emerged in the south of the country from a small fishing village in El Oro Province, and it is suspected that a fisherman introduced the index case was traveling north from Perú (Dixon et al. 2006). From 1991 to 2004 over 90,000 cases of cholera were reported in Ecuador, with most cases from coastal provinces. El Oro and Guayas provinces, located in southern coastal Ecuador, collectively represented one of two disease epicenters in the country. Recent studies indicate a high risk of a second epidemic in Ecuador due to the presence of important risk factors including the growth of vulnerable urban populations, decreased investment in cholera surveillance and prevention programs, increased flood risk associated with climate change, and a street food culture that includes eating raw shellfish (ceviche) (Malavade et al. 2011). In addition, Guayaquil (Guayas Province), the largest city in Ecuador, has been identified as the third most vulnerable city in the world to future flood risk (Hallegatte et al. 2013). Furthermore, it has been found that in populations with a high prevalence of blood group O, such as in Latin America, illness from cholera is more severe, and the requirements for rehydration and hospitalization of infected individuals are considerably higher (Swerdlow et al. 1994; Nelson et al. 2009). Given these conditions, there is compelling evidence that people in southern coast of Ecuador are a high-risk population and there is a critical need for active cholera surveillance in this region.

To address this, we evaluated local variability in the presence of *V. cholerae* in the estuarine environment surrounding the city of Machala, El Oro province, a site identified as a current and future coastal cholera hotspot (Escobar et al. 2015). We selected five sampling sites associated with estuarine water access in Machala, Ecuador, representing a range of economic and human activity conditions, in addition to different proximity to the ocean. Using water sampling methodology, coupled with laboratory identification of *V. cholerae* bacteria, we assessed the local environmental and pathogenic conditions over a period of ten months. Strengthening climate and water-sensitive infectious disease surveillance systems (WHO 2003; Zuckerman et al. 2007) and further understanding of the role of environmental factors in disease outbreak and transmission over time and space (Sedas 2007; Akanda et al. 2013) are urgently needed to target cholera and other climate and water sensitive diseases.

## Methodology

### Study Site

Machala is a port city of approximately 250,000 inhabitants, with major economic activities stemming from agriculture (bananas), aquaculture (shrimp farming), and fishing/shellfish collection, both small-scale and semi-industrial scale. Five sampling sites (Isla Jambelí, Boca del Macho, Puerto Bolívar Boca, Puerto Bolívar Adentro, and Héroes de Jambelí) were established within the Machala estuarine system (Figure 1), selected for maximum heterogeneity, to include highly built urban areas, ports, mangrove, and coastal areas. Isla Jambelí is on the outer edge of the coastal draining estuary, and the entrance to Jambelí is interspersed with mangroves and shrimp farms. Boca del Macho is the open edge of the inner estuary, in open water on shallow sand, with mangroves. Puerto Bolívar Boca is near the mouth of the open harbor, characterized by heavy boat traffic, commercial fishing, and residences lining the waterway, with mangroves and shrimp farming on the far side of the waterway. Puerto Bolívar Adentro is within the city, on the estuary, in an area characterized by residential low-income housing, with shrimp farms and mangroves across the Héroes waterway. Heroes de Jambelí is the most inland site, characterized by low income and poor quality housing built along mangroves at the edge of the city; outflow from the houses visibly drains directly into the water (Figure 1). The port city of Machala is an important sentinel surveillance site, due to its location along the Pan American highway, approximately 80km north of the Peruvian border, facilitating significant movement of people and potential pathogens by land and sea.

**Figure 1:**
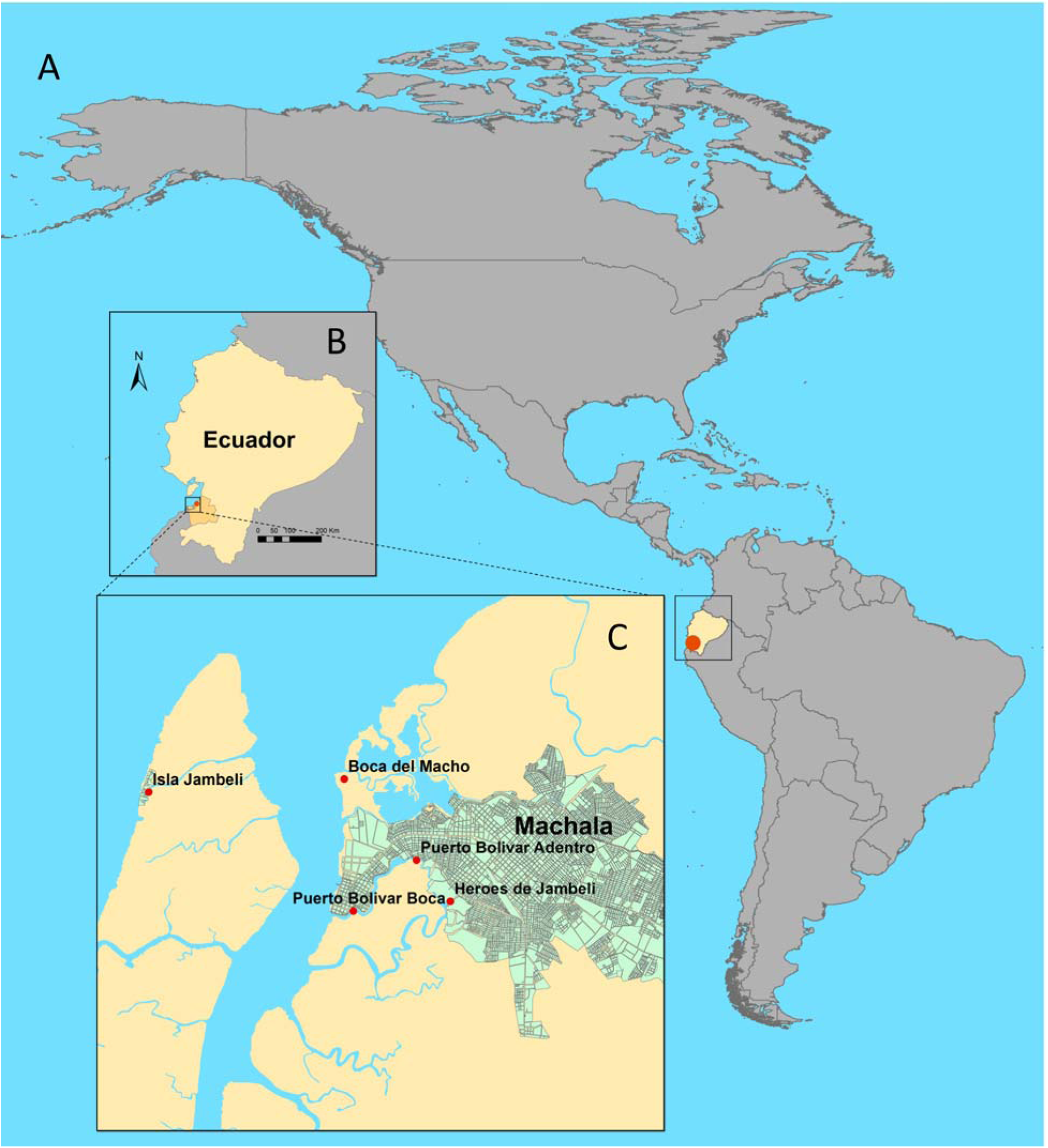
Location of sampling sites. **A.** Ecuador (in yellow) in South America, indicating the location of Machala (red point); **B.** Location of Machala on the southern coast of Ecuador (red point); **C.** Location of the five sampling sites: Isla Jambelí, Boca del Macho, Puerto Bolívar Boca, Puerto Bolívar Adentro, and Héroes de Jambeli (red points), in and around Machala (green).

### Collection of water samples and characteristics

At each of the five study sites (Figure 1), water sampling was conducted at high tide, twice monthly along a transect with 3 sub-sites spaced 250 m apart, and 3 replicates per subsite. Three 1L surface water samples per subsite were collected in sterile polypropylene bottles, and placed in coolers with ice for transport for the laboratory. We sampled biological, chemical, and physical water characteristics using a YSI water probe* (600 XLM V2 Sonde). We recorded Surface Temperature (°C), Conductivity, pH, Salinity, and Optic-T BGA PE (Phycoerythrin) [blue-green algae] (cells/ml, which we converted to cells/μl for ease of visualization), at each end of the transect.

### Laboratory analyses

Water samples were transferred to the laboratory in coolers for *V. cholerae* testing and were processed within 24 hours of collection. For laboratory analysis, a 1L water sample was filtered through a Whatman membrane No. 1 and 0.22 μm membrane (Millipore) by vacuum. Then, 10mL of Phosphate Buffered Saline (PBS) (pH 7.4) was pipetted onto the retained contents on the membrane and gently washed by pipette 15x. The PBS was left on the membrane to incubate at room temperature for 15 minutes prior to collection in 50mL conical tube. 3mL of membrane-washed PBS was enriched in 27mL alkaline peptone water (APW) (1% peptone, 1% NaCl, pH 8.6) and incubated for 24 hours at room temperature. Ambient room temperature in the laboratory was recorded daily. 5 mL of bacteria enriched with APW was then centrifuged at 4500 rpm for 10 minutes at 4°C, the supernatant was decanted, and the pellet was frozen at −80°C for DNA isolation.

### DNA isolation and PCR

Genomic DNA was extracted from bacterial pellet of the previous step with a QIAamp DNA mini kit (Qiagen), following manufacture instructions. Multiplex PCR was used for identification of cholera serogroups and the detection of toxigenic genes. Table 1 describes primers sets used to amplify the *rfb* region of O1 and O139 serogroups and the toxin subunit A (ctxA) and toxin coregulated pilus (tcpA) genes. For both duplex PCRs, master mix was as follows: 0.05 U/ul of JumpStart REDTaq DNA Polymerase (Sigma), 1X buffer, 0.2 mM dNTPs, 0.2 mM of each primer set, 1 μl of template and ultrapure water to a final volume of 25 μl. The amplification program for diagnosis of serogroups was adapted from Hoshino et al. (1998), using the following conditions: 5 minutes at 94°C, 35 cycles of 94°C for 1 minute, 55°C for 1 minute and 72°C for 1 minute and final extension of 72°C for 7 minutes. Positive samples for either or both serogroups were subjected to toxigenic genes duplex PCR. The amplification program was according to conditions described in Kumar et al.(2010): 3 minutes at 94°C, 30 cycles of 94°C for 30 seconds, 59°C for 30 seconds and 72°C for 1.2 minutes, and final extension of 72°C for 10 minutes. To validate our multiplex PCR methods, the PCR products were resolved in a 2% agarose gel and sequenced to verify gene amplification.

**Table 1.**
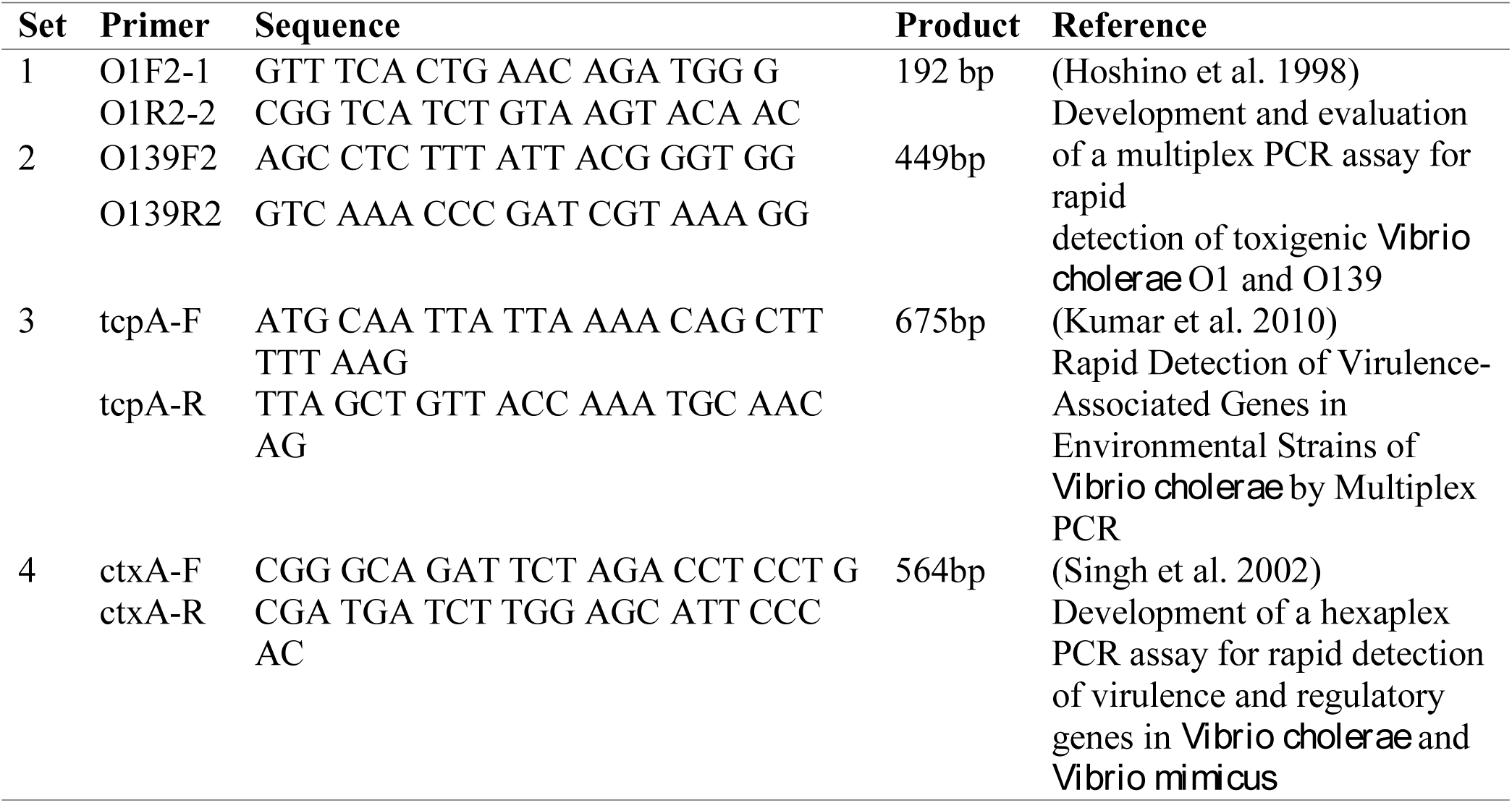
PCR primers set used in this study

### Statistical analyses

As the data were not normally distributed, we conducted non-parametric tests throughout. We characterized the sampling sites for each environmental variable: Temperature, pH, Salinity, and BGA, conducting Kruskal-Wallis rank sum tests on site means, and on monthly means. We then examined whether *V. cholerae* prevalence at sites, and strain (i.e., O1, O139) prevalence separately, was associated with environmental variables using a series of non-parametric Kendall’s tau correlations.

## Results

### Water sample characteristics

The probe recorded a range of 9-104 readings at each sub-site biweekly for 10 months. We pooled all readings by month for analyses. Our sites differed significantly in water characteristics (Figure 2), as shown by a series of Kruskal-Wallis rank sum tests (Temperature: χ^2^= 206.19, df = 4, *p* < 0.0001; Salinity: χ^2^= 2257.5, df = 4, *p* < 0.0001; pH: χ^2^= 1347.3, df = 4, *p* < 0.0001; BGA χ^2^= 1824.8, df = 4, *p* < 0.0001). We found that Héroes de Jambelí, the most inland site, had the highest recorded BGA, and that Isla de Jambelí, the most coastal site, had the highest recorded salinity; while there were statistical differences between all sites in all characteristics, there were no clear outliers in pH or temperature. Our sites exhibited significant change in water characteristics across months (Figure 3), as shown by a series of Kruskal-Wallis rank sum tests (Table 2). Temperature was lowest in August for all sites – likely reflecting pacific upwelling, which cools the water, regardless of air temperature. Salinity at the most inland site, Héroes de Jambelí, was consistently lowest, and showed the smallest change across months, while the other sites had a decrease in salinity in May, then a rise from July-December. Isla de Jambelí had the highest salinity, reflecting its location on the most coastal site. BGA was highest at the most inland site, Héroes de Jambelí, peaking in May, lagging temperature by a month. BGA shows the least temporal or spatial clustered pattern and has no obvious seasonality across the year. Heroes de Jambeli, however, registered the highest BGA values during the study (∼25,000). pH appears to peak in December-January across all sites, with a decrease in July-August; the coastal and inland sites showed low pH values across seasons, while Boca del Macho registered consistently high pH values across months.

**Table 2.**
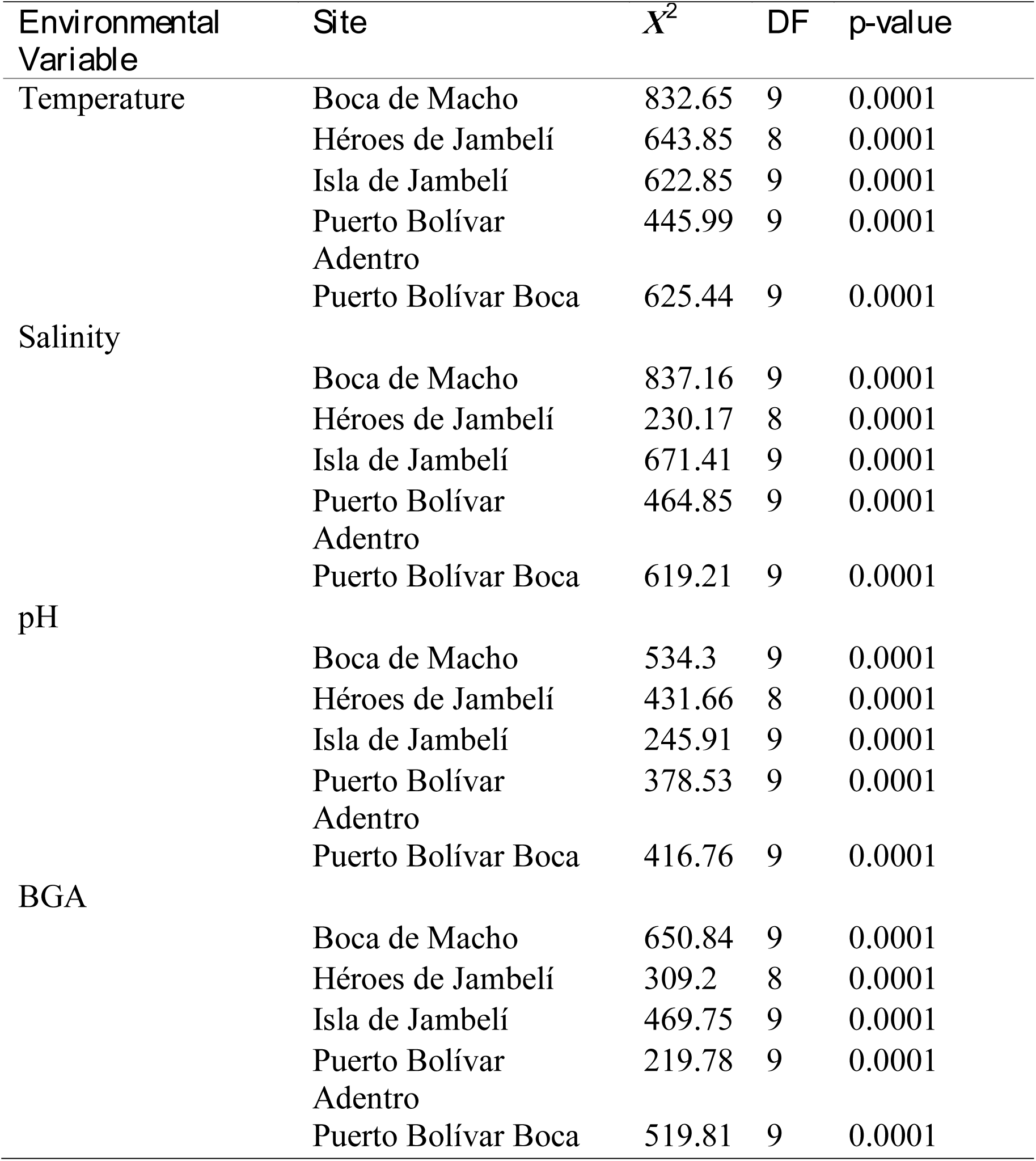
Kruskal-Wallis rank sum test results for each site and environmental variable differences by month.

**Figure 3:**
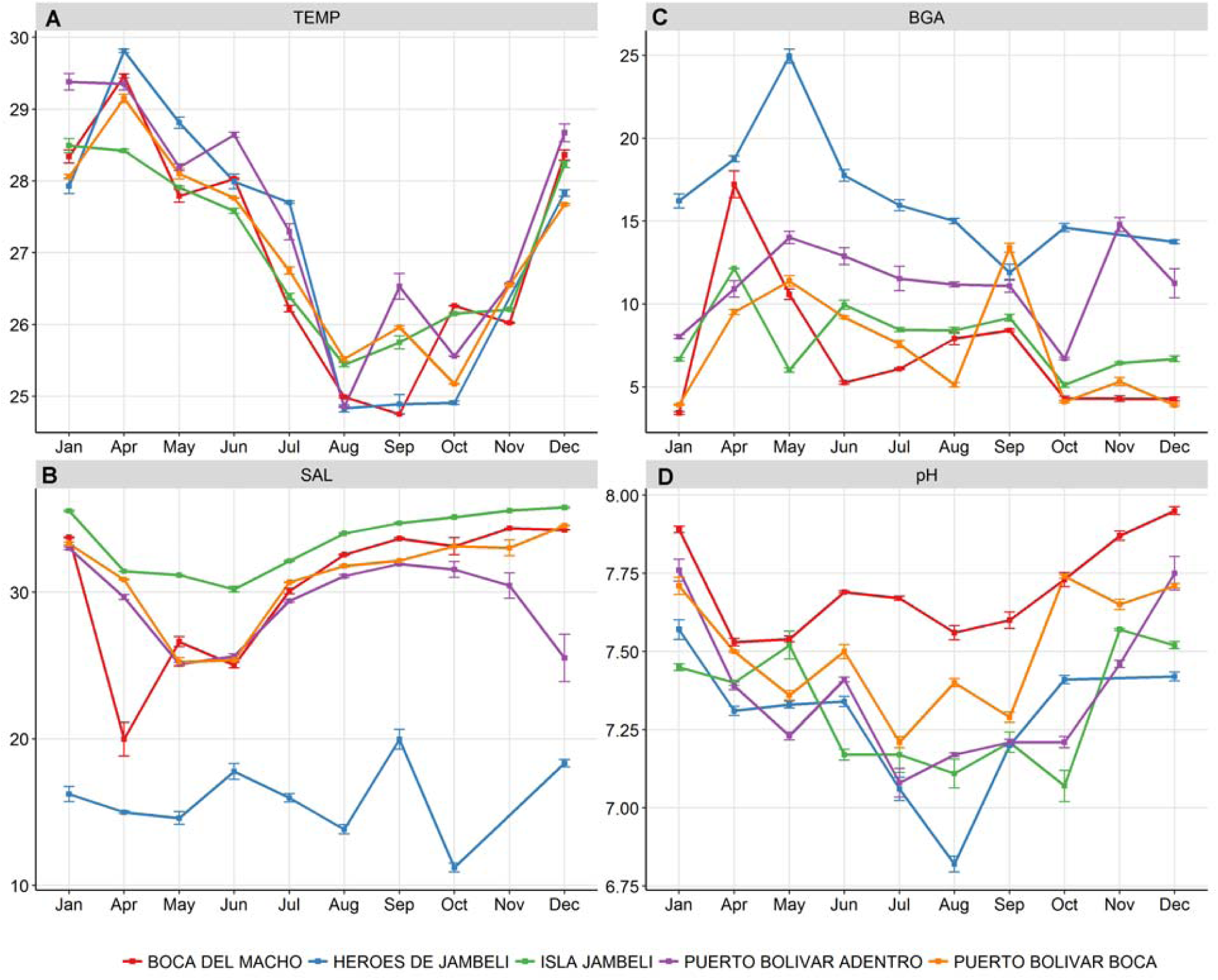
Environmental features in the study period. Water characteristics by month (means and standard errors) and sites: **A**. Temperature (TEMP, °C), **B.** Salinity (SAL), **C.** measured total concentration of blue-green algae (BGA, cells/μL), and **D.** pH.

**Figure 2:**
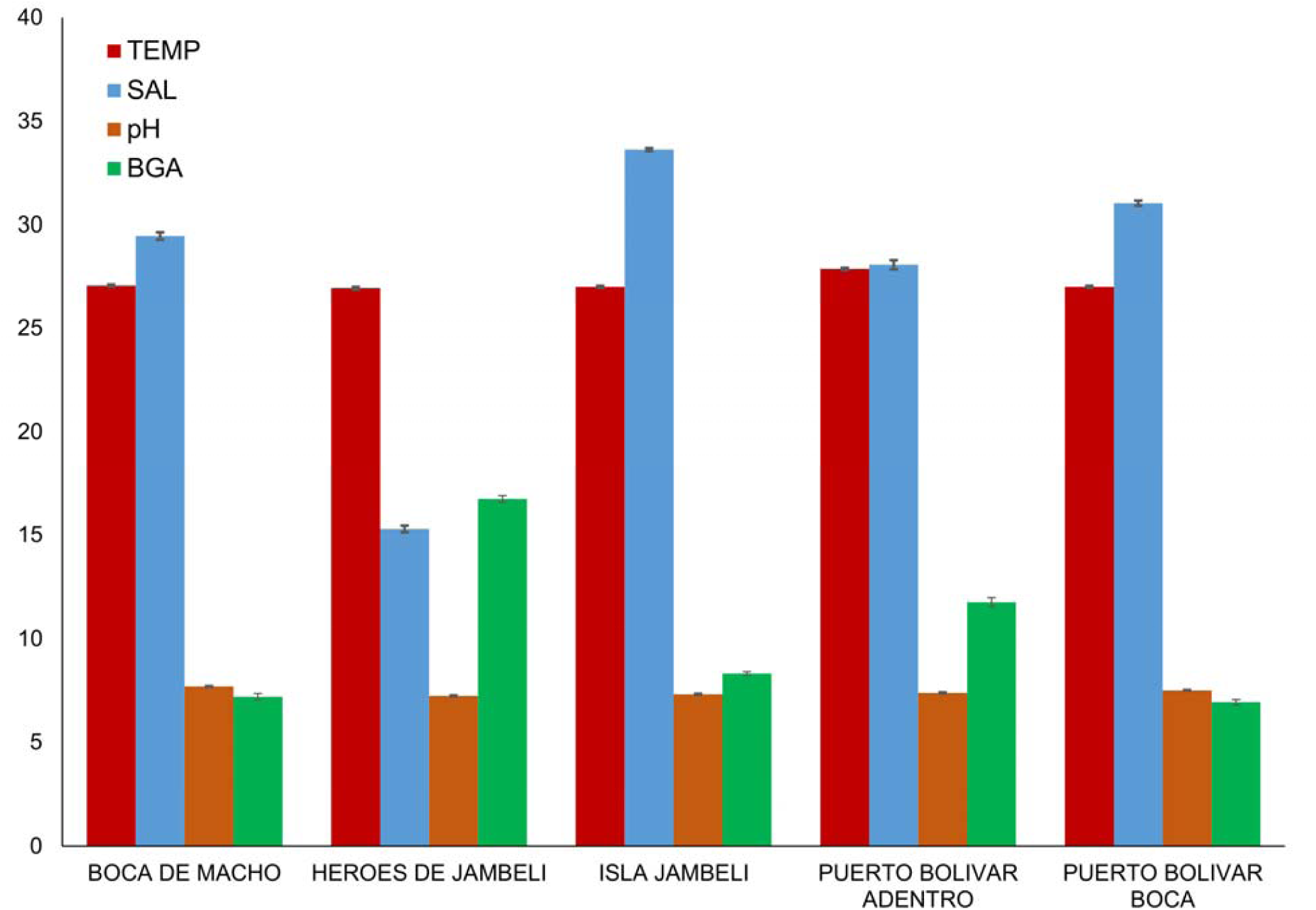
Environmental features in the study area. Water characteristics by site (means and standard errors Temperature (TEMP, °C), Salinity (SAL), pH, and measured total concentration of blue-green algae (BGA, cells/μL).

### Laboratory analyses

Of a total of 405 individual water samples, collected between May – September, 382 were diagnosed by multiplex PCR. We found 139 (36.4%) samples positive for *V. cholerae*, and 243 (64%) negative. We found both O1 and O139 serogroups of *V. cholerae* present in the estuarine system studied in Machala, Ecuador. Serogroup O139 was predominant; 118 (83.5%) samples were O139, and 51 (35.3%) were O1 (30 samples contained both). We were able to detect *V. cholerae* during each of the 5 months sampled, nevertheless we found that prevalence decreased drastically in July (Figure 4). By sequencing the samples, we confirmed that the PCR protocol applied was appropriate for detection of *V. cholerae* serogroups O1 and O139 strains. Results from the toxin subunit A (ctxA) and toxin coregulated pilus (tcpA) analyses were all negative. We found no evidence for persistent environemental toxigenic *V. cholerae* in our samples.

**figure 4:**
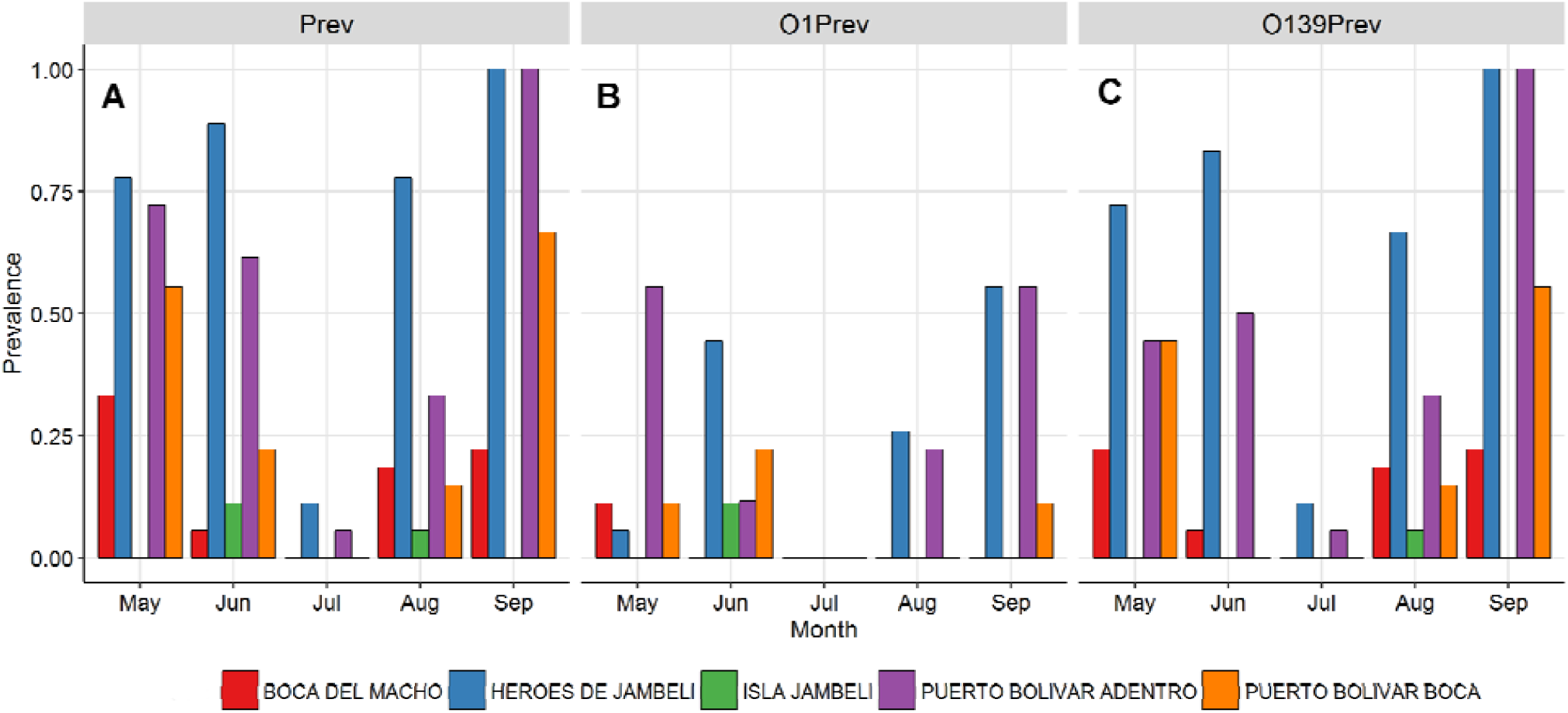
*Vibrio cholerae* detection. Monthly site prevalence of **A.** cholera as given by positive PCR test, **B.** O1 strain, **C.** O139 strain.

### *Vibrio cholerae* prevalence

We pooled water samples within sites, to derive monthly *V. cholerae* prevalences across and within sites (prevalence = positive/total samples tested). Overall monthly prevalence of *V. cholerae* ranged from 0.3 (n=68) in July to 0.58 (n=45) in September, with site prevalence ranging 0-1, with a mean monthly site prevalence of 0.35 (Figure 4A). Individual strain prevalence was generally higher for O139 than O1, but we see that Puerto Bolívar Adentro and Héroes de Jambelí were *V. cholerae* positive in every month and also had higher prevalences than the other sites (Figure 4B and 4C). We found that the prevalence of *V. cholerae*, and O139 and O1 strains separately, were significantly associated with higher BGA (blue-green algae densities), and that prevalence of *V. cholerae* the O1 strain were significantly associated with lower salinity. We found no significant association between prevalence and temperature or pH (Table 3).

**Table 3:** Kendall tau tests for correlation between prevalence of cholera, and each strain separately, and environmental variables at sites, pooled monthly.

| Environmental Variable | Prevalence | Kendall's $\tau$ | z | p-value |
| --- | --- | --- | --- | --- |
| Temperature | *V. cholerae* | 0.08 | 0.56 | 0.57 |
|  | O1 | 0.12 | 0.80 | 0.42 |
|  | O139 | 0.02 | 0.12 | 0.91 |
| Salinity | *V. cholerae* | -0.34 | -2.35 | **0.02** |
|  | O1 | -0.35 | -2.29 | **0.02** |
|  | O139 | -0.28 | -1.92 | 0.06 |
| pH | *V. cholerae* | -0.10 | -0.68 | 0.49 |
|  | O1 | -0.14 | -0.92 | 0.36 |
|  | O139 | -0.10 | -0.64 | 0.52 |
| BGA | *V. cholerae* | 0.55 | 3.76 | **0.00** |
|  | O1 | 0.48 | 3.09 | **0.00** |
|  | O139 | 0.49 | 3.34 | **0.00** |

### Discussion and Recommendations

We found evidence of an environmental reservoir of *V. cholerae* in the estuarine waters of Machala, Ecuador, in 2014. We confirmed the presence of *V. cholerae*, including pandemic strains O1 and O139. We cannot rule out ongoing toxigenic presence, but we did not detect it in our samples. Prior to 1961, epidemics of cholera were associated only with O1 strain, both classic, and later, El Tor type, with the pathogenic O139 strain appearing in the 1992 pandemic in the Bay of Bengal, arising from genetic exchange with O1 El Tor (Lipp et al. 2002). Other pathogenic O1 strains are thought to have evolved and emerged independently, such as the U.S. Gulf Coast O1 strain, and the Australian clone (Faruque et al. 1997), and there is evidence that toxigenic strains can arise from nontoxigenic environmental strains (Jiang et al. 2000), including toxigenic O139 strain. Without further investigation, we cannot definitively attribute the ongoing environmental *V. cholerae* presence to a residual persistent reservoir since 1992, or whether introduction of strains has repeatedly occurred from ballast water exchange at the port. However, we see persistence of the two strains most commonly associated with epidemics around the globe.

Our sites exhibited considerable seasonal and spatial heterogeneity in water characteristics and *V. cholerae* prevalence, with clear peaks (and troughs) during specific months. For example, we found peak *V. cholerae* prevalence in September, with highest values in two sites: Héroes de Jambelí and Puerto Bolívar Adentro (Figure 1). These sites are characterized by low income housing on the edge of the city, while being inland sites, facing mangroves and shrimp farms, and were found to have *V. cholerae* present in every month sampled. The lowest *V. cholerae* prevalence was found in July, in which only the two most inland sites had detectable *V. cholerae*. Water temperature had the clearest temporal pattern, falling rapidly through July, likely corresponding to Pacific upwelling, cooling the waters, and increasing nutrients in the system (Strutton et al. 2001). Unsurprisingly, we found the lowest salinity in the most inland site, Héroes de Jambelí, with a higher concentration of BGA than in other sites. This is in contrast to Isla Jambelí, a small island community furthest from the mainland and closest to the ocean, with high salinity due to its coastal location; however, it did not have significantly lower BGA than other sites.

We found that the timing of *V. cholerae* prevalence was coupled to the water characteristics we measured. For example, temperature, BGA, and pH decreased in most sites through July/August, as did the prevalence of *V. cholerae*, but we only demonstrated statistically significant associations between prevalence and site and month specific salinity and BGA. Average ocean salinity is around 35 ppt, while freshwater rivers average around 0.5 ppt; this estuarine system is a mixed or brackish system, ranging from the lower average of around 15 ppt at our most inland site, to a high approaching 34 ppt at our coastal site. Our most inland site, represents optimal salinity for *V. cholerae* growth (Montilla et al. 1996). We detected *V. cholerae* at a range of salinities, finding a negative correlation with increasing salinity, suggesting that lower salinity permits a suitable environment for the growth of *V. cholerae*, but higher salinities approaching oceanic concentrations do not completely prohibit growth. This finding is consistent with previous work demonstrating the suitability of coastal oceans for *V. cholerae* (Strutton et al. 2001), but reveals a finer scale relationship with salinity in an estuarine system, up the gradient to fresh water.

BGA (blue-green algae; a.k.a. cyanobacteria) are photosynthetic prokaryotes that can be found in freshwater, marine, and terrestrial environments (Stanier and Bazine 1977). The photosynthetic pigments of cyanobacteria include chlorophyll-*a* and the phycobiliprotein phycoerythrin. Here we used BGA values to characterize water features and because BGA has previously been associated with *V. cholerae* persistence (Epstein 1993). Temperature increase, coupled with high nutrient load, low flow, and thermal stratification, generally results in increased growth rates of cyanobacteria, and its dominance in the phytoplankton community (Davis et al. 2009; Elliott 2010; Huber et al. 2012). This could explain the high BGA values early in the year (Figure 3). In addition, warm temperatures promote increases in the number of days where BGA biomass exceeds warning thresholds established by WHO (Davis et al. 2009; Elliott 2012). High temperature also influences water column stability and mixing depth, producing favorable conditions for BGA blooms (Robarts and Zohary 1987; Stal et al. 2003). This association of temperature increase with BGA blooms is consistent across coastal, estuarine, and inland waters (Paerl 1988), illustrating the suitability of tropical estuarine waters for environmental *V. cholerae* growth and persistence. We recommend that long term monitoring to measure BGA biomass should be considered at minimum in Heroes de Jambeli and Puerto Bolivar, the sites reporting the highest BGA values (Figure 2). This is particularly important, looking to the future, considering that a rise in water temperature – which we expect with global climate change - is associated with BGA emergence (Wasmund 1997; Kanoshina et al. 2003; Suikkanen et al. 2007).

This study was conducted in an average climate year, providing a preliminary framework for monitoring coupled *V. cholerae* – estuarine dynamics for potential emergence of cholera outbreaks in the region. We recognize that this is a preliminary study, and with limited sample size, precluding larger conclusions or significant predictive power for the future. As the first study of this type in the region, and as a prototype for small-scale epidemic surveillance platforms, we provide evidence of the presence of these persistent strains, and the capacity to conduct such monitoring. This is particularly useful baseline information for anticipating El Niño years, extreme climate events associated with warming temperatures of surface ocean water and increased rainfall and flooding events. The year following the study, an El Niño year, saw severe flooding throughout Machala. Climate change projections indicate that the frequency of extreme El Niño events will increase in the future (Cai et al. 2014), increasing the risk of water-borne diseases endemic in the region, such as cholera, typhoid, and leptospirosis. This study provides valuable information, and our recommendation is to add *V. cholerae* back into the public health agenda, to consider infectious diseases beyond the already important vector-borne diseases, such as dengue fever, chikungunya, zika, and malaria.

Indeed, in May 2016, two years after the initiation of this study, a case of cholera was reported in Machala, after approximately 12 years with no case reports in Ecuador (2016). An immuno-compromised individual was confirmed positive for *V. cholerae* serotype O1 nontoxigenic, by the National Public Health Research Institute of the Ministry of Health. Our team diagnosed the patient using the same PCR assay described here, because we were the only lab in the region with this capacity. Although the source of the infection was not confirmed, this case report suggests a worrisome link to environmental transmission, underscoring the importance of our results.

This study highlights the urgency for active epidemiological surveys and the need for public health interventions to reduce the risk of water-borne pathogens in vulnerable populations from a holistic social-ecological perspective. The community Héroes de Jambelí is a low-income peri-urban settlement with less than 50 families, established informally in 2002. The community continues to lack adequate access to piped water, sewerage, and garbage collection due to their status as an illegal settlement. Simple bamboo homes have built over the mangrove system, with direct discharge of wastewater into the estuary. At the same time, this community’s livelihood depends on artisanal fisheries (e.g., crabs, mollusks) from these same estuaries. This vulnerable coupled human-natural system results in high risk of emerging epidemics from water-borne pathogens.

## Acknowledgments

We would like to thank the citizens and authorities of Machala, Ecuador, for their continued support of our research infectious diseases epidemiology and monitoring. We thank Carlos Enriquez for his hard work in the field.

